# Automatic local resolution-based sharpening of cryo-EM maps

**DOI:** 10.1101/433284

**Authors:** Erney Ramírez-Aportela, Jose Luis Vilas, Roberto Melero, Pablo Conesa, Marta Martínez, David Maluenda, Javier Mota, Amaya Jiménez, Javier Vargas, Roberto Marabini, Jose Maria Carazo, Carlos Oscar S. Sorzano

## Abstract

Recent technological advances and computational developments, have allowed the reconstruction of cryo-EM maps at near-atomic resolution structures. Cryo-EM maps benefit significantly of a “postprocessing” step, normally referred to as “sharpening”, that tends to increase signal at medium/high resolution. Here, we propose a new method for local sharpening of volumes generated by cryo-EM. The algorithm (LocalDeblur) is based on a local resolution-guided Wiener restoration approach, does not need any prior atomic model and it avoids artificial structure 1 factor corrections. LocalDeblur is fully automatic and parameter free. We show that the new method significantly and quantitatively improving map quality and interpretability, especially in cases of broad local resolution changes (as is often the case of membrane proteins).

## Main text

Cryo-Electron Microscopy (cryo-EM) is engaged, on the methodological front, to constantly improve the quality of its reconstructed maps, so as the subsequent construction of atomic models become more accurate. Addressing this problem, the so-called sharpening methods have been recently introduced in cryo-EM workflows as a post-processing step^1,2^, after being in common use in X-ray crystallography for a much longer time^3,4^. In cryo-EM the most widely spread method so far is a structure factor modification based on the Guinier plot, also known as B-factor correction^2^. The general idea behind this approach is to overcome the loss of contrast at high resolution by boosting the amplitudes of structures factors in that resolution range while at the same taking into account some form of measure of the frequential signal to noise ratio, this latter being a correction in order to limit noise amplification. The B-factor is the slope of the amplitudes falloff which will be boosted. In practice, however, the results are very similar to applying a global B-factor flattening at most frequencies, as it will be clear in the example we will analyze.

Sharpening algorithms can coarsely be classified as global and local. However, up to our knowledge, all of them make use of the basic amplitude correction method introduced above. Thus, global sharpening determines a B-factor value, which is then considered for the whole map correction. Relion post-processing^5^ belongs to this group, directly working on the Guinier plot, as newer methods also do, such as AutoSharpen in Phenix Package^6^, which tries to optimize the B-factor based on a merit function that maximizes the connectivity at the same time that minimizes the surface of the resultant sharpened map. Global approaches apply the same transformation to the whole map, neglecting the fact that different regions might present different resolutions. In contrast, in the local sharpening group, we comment on LocScale^7^. It compares the radial average of the structure factors inside both a moving window of the experimental map and of the map calculated from the corresponding atomic model. Then, it locally scales the map amplitudes in Fourier space to be in agreement with the atomic model. An obvious limitation of this method, of course, is the essential requirement for a starting atomic model, which may restrict its applicability.

To overcome the limitations presented above, a fully automatic and parameter-free local sharpening method based on local resolution estimation is presented in this work. The algorithm, named LocalDeblur, acts as a local deblurring. Our method solves the sharpening problem using a local gradient descent version of the Wiener filter. The root of this algorithm is a space-variant filter. We consider that the observed cryo-EM density map has been obtained by the convolution with a local, lowpass filter whose frequency cutoff is given by the local resolution estimate. We use this information to compute the sharpened map by an iterative steepest descent method. LocalDeblur does not require an atomic model and, consequently, is bias free in this regard. Full derivation of LocalDeblur is presented in Methods section.

## Results

The performance of LocalDeblur was tested with three different unsharpened cryo-EM density maps from the recent EMDB map challenge (JSB 2018, Special Issue): TRPV1 channel (EMD-5778)^8^, *Plasmodium falciparum* 80S ribosome (EMD-2660)^9^ and *Thermoplasma acidophilum* 20S proteasome (EMD-6287)^10^. The local resolution needed for our algorithm was calculated using MonoRes^11^ (other algorithms could be used).

Naturally, all current methods for cryo-EM sharpening considerably enhance map interpretability; consequently, and in order to make a clear case for the significant step forward in the field represented by an automatic and parameter-free method such as the one presented in this work, we performed comparisons between LocalDeblur and current versions of the main sharpening methods in use in the field, such as Relion post-processing^5^, Phenix AutoSharpen^6^ and LocScale^7^. Two sets of comparisons were performed, first analyzing the maps (Fig. 1 and Fig. 2), and then analyzing their corresponding Guinier plots (Fig. 3 and Supplementary Fig. 2).

**Figure 1.**
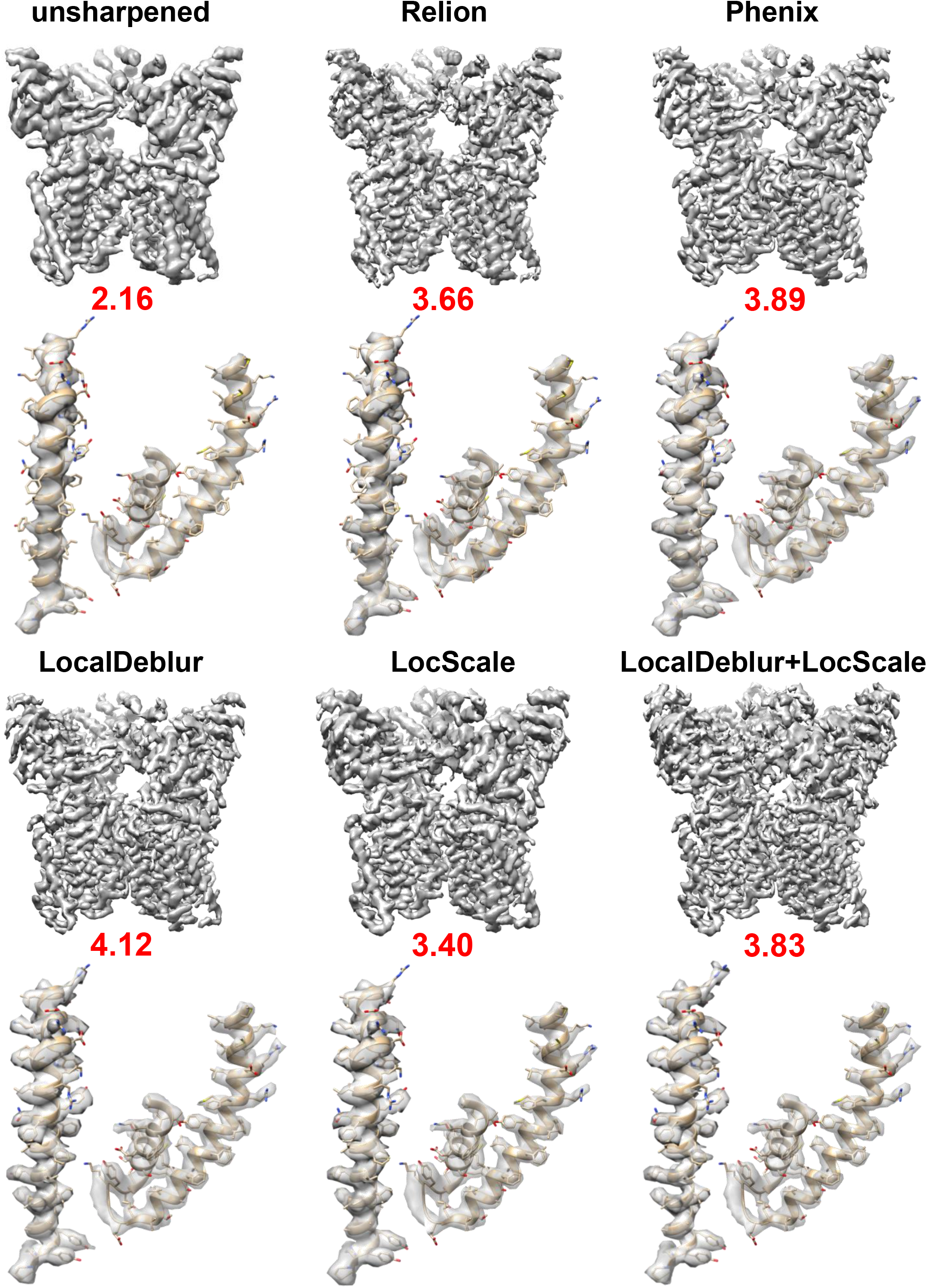
Sharpened map of TRPV1 (EMD-5778) generated with LocalDeblur and comparison with current main sharpening methods (Relion post-processing, Phenix AutoSharpen and LocScale). Below each map, the value of EMRinger score is shown and the densities corresponding to 419–456 (left) and 568–642 (right) residues are represented.

**Figure 2.**
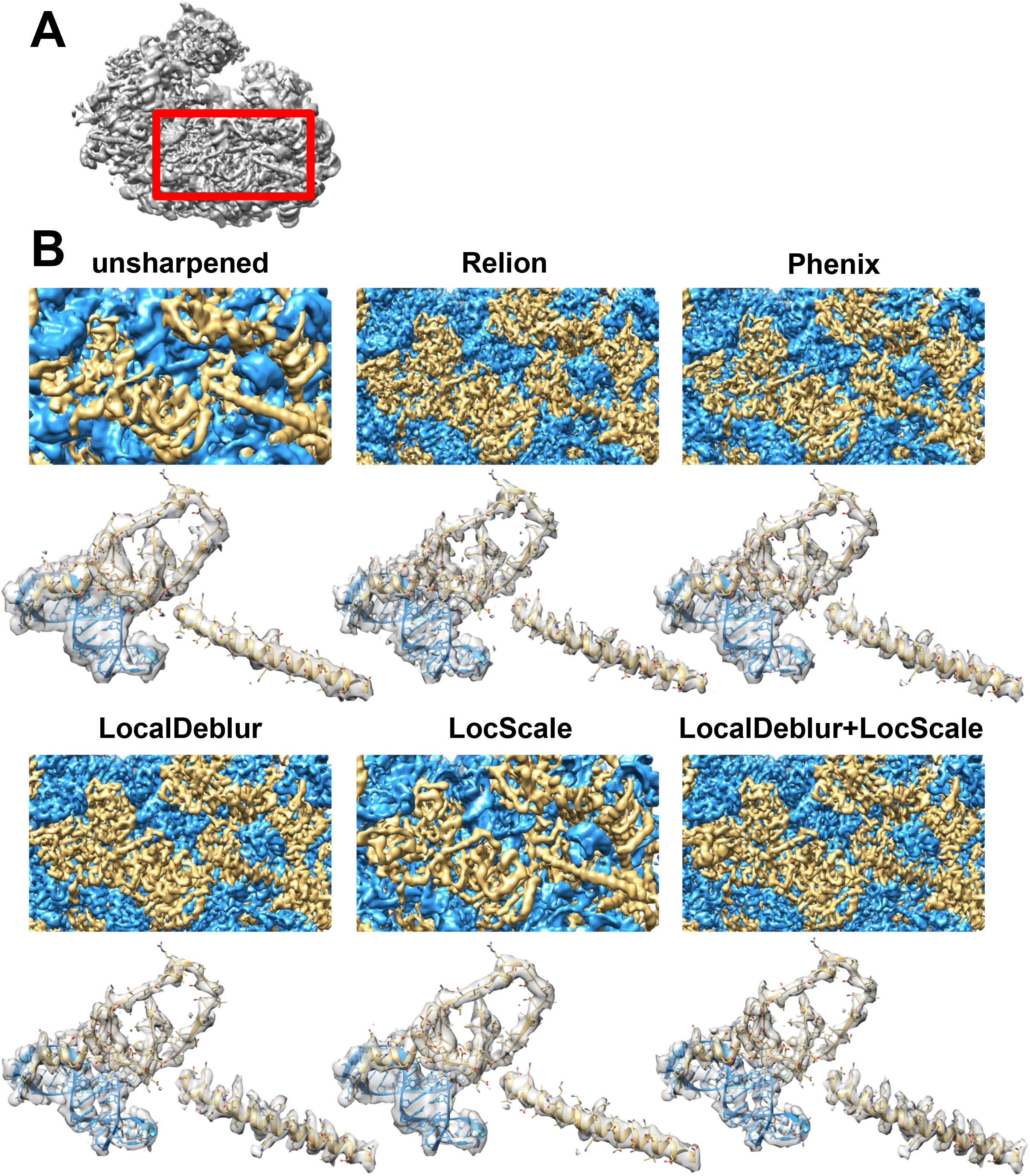
Sharpened maps of *Plasmodium falciparum* 80S ribosome (EMD-2660). (A) The whole density map from 80S ribosome is shown. The red frame corresponds to the density enlarged in panel B. (B) Sharpened map of 80S ribosome generated with LocalDeblur and comparison with the main sharpening methods (Relion postprocessing, Phenix AutoSharpen and LocScale). Only the section corresponding to the red frame in A is shown. The RNA density is represented in blue and the amino acid density in yellow. Below each sharpened map, the density for 149–186 (chain-K) (left), and 3712–3727, 3761–3775 (chain-A) and 283–381 (chain-E) (right) residues are represented.

**Figure 3.**
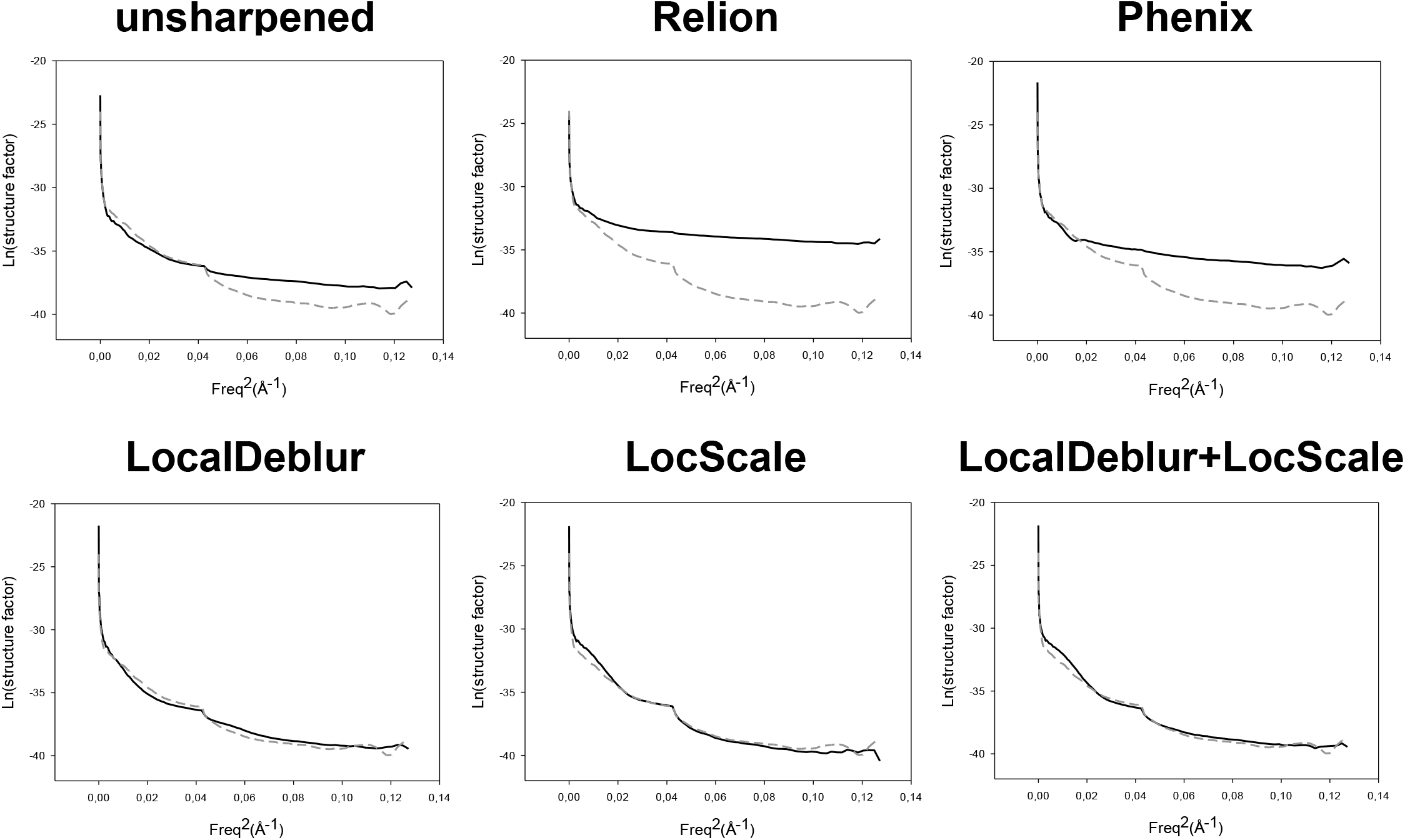
Guinier plots for each sharpened map represented in Figure 1 are shown. The profile corresponding to the density map generated from the atomic model (PDB ID: 3j5p) is superimposed as a dashed line representing our “target result”. Note how profiles obtained by LocalDeblur are very similar to the target ones.

## Comparison of sharpening methods

Qualitatively, it can be clearly appreciated in Figs. 1 and 3 that all sharpened maps have more details than the original maps, but that in all LocalDeblur maps the α-helical pitch is always more clearly delineated and the density for most side chains is considerably better defined than in the other methods, which can substantially facilitate the construction of the corresponding atomic model. The application to TRPV1 (Fig. 1) demonstrates LocalDeblur usefulness for membrane proteins, while 80S Ribosome maps (Fig. 2) show that LocalDeblur works very well with macromolecules that contain both amino acids and nucleotides. Indeed, LocalDeblur improves the contrast between both and allows a better definition of high-resolution features.

Next, we combined LocalDeblur and LocScale. The objective was to check if once LocalDeblur has increased the interpretability of the map (without any additional information), then, the knowledge of an initial atomic model can slightly enhance the sharpened map with LocScale. Figs. 1–3 and Supplementary Figs. 2 present the results of this combination. As shown, if the atomic model of the EM map is available, a combination of LocalDeblur and LocScale indeed produce a slightly better sharpened map (this seems to be case dependent, being much clearer for the ribosome than for TRPV1), but the small increase in interpretability comes at a very high price, in that an initial structural model has to be provided in LocScale, while this is not required in LocalDeblur.

### Map quality determine quality of atomic model

We further quantitatively studied the influence of the each of the sharpening methods compared in this work on the quality of the fitting of atomic models for the specific case of TRPV1. In this way, we refined the atomic model of TRPV1 (PDB ID: 3j5p) for each sharpened density map. The initial atomic model was fitted into the sharpened maps of TRPV1 using UCSF Chimera^12^. Subsequently, the fitted model was further refined using Coot^13^ and, then, underwent 5 iterations of real-space refinement in Phenix^14^, including rigid-body, model morphing, local real-space fitting, global gradient-driven and simulated annealing refinement. At the end of the refinement, the quality of the geometrical parameters in every case was improved with respect to the ones of original model. For quantitative comparisons, EMRinger^15^ values are shown under each map in Fig. 1, while other quality measures determined using MolProbity^16^ are presented in Supplementary Table 1. The highest EMRinger score is achieved for the refined atomic model with LocalDeblur map, indicating that amino-acid side chains are very chemically realistic and fit very well into the density map.

## Comparison of Guinier Plots

The second set of comparisons addresses the behavior of structure factors as maps are sharpened by the different methods. In Fig. 3 we present the data corresponding to TRPV1 and in Supplementary Fig. 1 those for the 80 Ribosome, showing total coincidence. Focusing on TRPV1, the Guinier plot of the radially-averaged profile of each sharpened map is shown, where the profile corresponding to the density map generated from the atomic model (PDB ID: 3j5p) is plotted with 6 a dashed line. It is readily noticed that this latter profile shows a decay and that it has a clear peak at ∼4.9 Å, which is within the range of characteristic distances of secondary structure elements (7 to 4 Å)^17^. However, in maps obtained with Relion post-processing and Phenix AutoSharpen, the characteristic features of the original structure factors at middle and high resolution are lost, including their frequency decay and the peak mentioned above. On the other hand, methods that take into account local characteristics of the maps (such as LocScale and LocalDeblur) are much more consistent with the expected structural factors corresponding to the structure under investigation. In general, local sharpening methods reproduce overall radial structure factor profiles much better than methods based on global B-factor. This fact correlates very well with the previous observation that secondary structure elements are substantially better presented in LocalDeblur than in any other method without prior information, even if the atomic model was not used at all in the calculations.

All these experiments point out into the direction that the application of global sharpening based on the B-factor value may not be an optimal procedure in cryo-EM. Indeed, owing to intrinsic characteristics of the macromolecules and errors during the reconstruction workflow^18^, different parts of the maps can have varied resolutions^11,19,20^ and, consequently, they may require different levels of sharpening/blurring for optimal interpretation. In these cases, the selection of an appropriate global sharpening/blurring B-factor value becomes an impossible task, since it does not take into account the local signal to noise ratio of the maps. A simple example is the proteasome core (EMD-6287) (Fig. 4), a “classic” example in cryo-EM of a stable specimen. Indeed, the local resolution map determined with MonoRes^11^ indicates a relatively narrow range of local resolution values at the center, but with very significant degradations especially in the distal peripheral regions. Clearly, even for this “stable” specimen, a global B-factor-based sharpening method is not adequate to correctly analyze the density map corresponding to these lower resolution regions (Fig. 4B for details).

**Figure 4.**
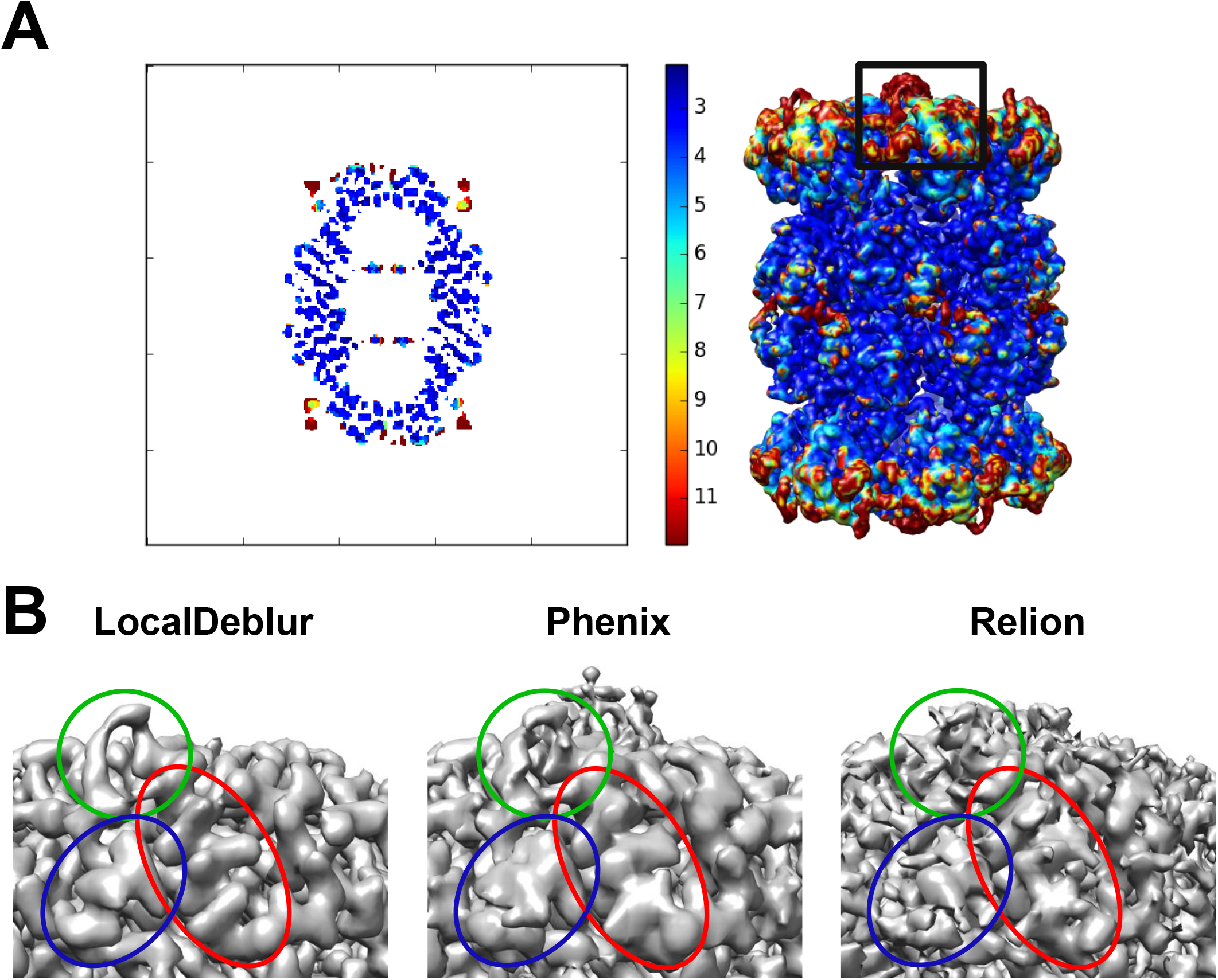
MonoRes results and Sharpened maps of *Thermoplasma acidophilum* 20S proteasome (EMD-6287). (A) MonoRes resolution slice and resolution map for the 20S proteasome. The black frame correspond to the density enlarged in panel B. (B) Close-up of the density maps generated with LocalDeblur, Phenix Autosharpen and Relion post-processing, respectively, displaying the peripheral proteasome fragment marked in (A). Note how LocalDeblur map is better defined in the indicated areas.

## Conclusions

In this paper we have introduced a new method to locally sharpen electron microscopy maps. The concept behind our algorithm is quite straight forward: Understanding local resolution as a local blurring of an otherwise accurate map called sharpened map, a fact that immediately establishes a simple convolution relation between the measured and sharpened maps via local resolution. The sharpened map is then obtained by deconvolution. Indeed, the new method is automatic, parameter free and only requires an estimate of the local resolution.

Results for different types of macromolecules have been used to test the algorithm. In particular, the test structures were carefully chosen to cover many scenarios as they are, membrane protein (TRPV1), high resolution volumes (proteasome), and maps with broad resolution ranges and distinct biochemical components (ribosome 80S). In all cases LocalDeblur has shown an excellent 8 performance in comparison with other methods, improving the interpretability of the maps and increasing the fitting quality of an atomic model.

Moreover, up to our knowledge, LocalDeblur is the first local sharpening approach based on local resolution, without the need to use a prior initial structural model and avoiding somehow artificial processes, as high frequencies boosting approaches based on B-factor quasi-flattening. In this sense, we have clearly shown that the new local method produces Guinier plots more similar to the Guinier plot of the corresponding atomic structures they were compared, totally outperforming global methods.

## Methods

### Implementation

The algorithm is publicly available from *Xmipp^21^* (http://xmipp.cnb.csic.es) and integrated in the image processing framework *Scipion 1.2.1^22^* (http://scipion.cnb.csic.es). In the near future is it expected to be offered as a standard Scipion Web Services Tools^23^, which would greatly facilitate the occasional use of the program without any local installation. LocalDeblur requires as input an unfiltered 3D reconstruction Cryo-EM density map and a local resolution map calculated either with MonoRes^11^ or ResMap^19^.

### Local deblurring method (LocalDeblur)

Consider two electron density maps ***v**_sh_* and ***v**_obs_* respectively called sharpened and observed, such that ***v**_obs_* is a degraded reconstruction map obtained from ***v**_sh_* as 
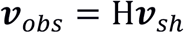

The volumes are represented by vectors in lexicographic order, and *H* is a blurring operator responsible for degrading the sharpened map. The definition of *H* is a local filter which acts by applying a filter bank upon the map ***v**_sh_*. Each filter in the bank is a bandpass filter centered at resolution *R_i_*, and represented by a matrix *H^i^*. Each channel in the filter bank is locally weighted according to the distance between the local resolution, ***R**_local_*, and the resolution of the filter. Note that ***R**_local_* is another vector of the same size as the input map, whose *j*-th entry is the estimate of the local resolution at the *j*-th location. We get these estimates of local resolution using MonoRes. The local weight is given by a diagonal matrix, *W^i^*, whose *j*-th entry is 
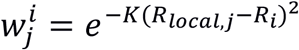

Empirically we have observed that casts good results for all tests we performed. This weight matrix tends to favor the contribution of voxels to the frequencies associated to their local resolution. Therefore, the matrix *H* can be mathematically written as 
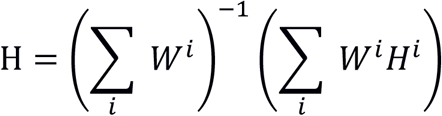

Note that the denominator should be introduced as a normalization term in the filter.

Our method solves the sharpening problem using a gradient descent version of the Wiener filter. For completeness, let us succinctly give here a short derivation of it. The Wiener filter can be regarded as the solution of a Bayesian restoration problem in which the noise and the signal both follow multivariate Gaussian distributions with zero means and covariance matrices Σ*_n_* and Σ*_x_*^24,25^. The Maximum a posteriori estimate of the signal would be given by 
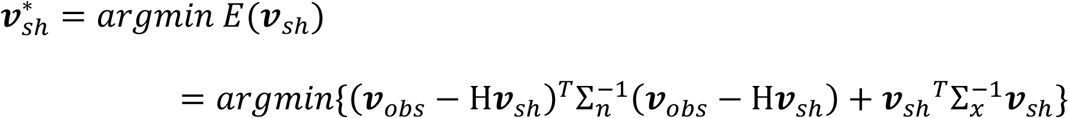

Note that if the signal covariance matrix is assumed to be of the form 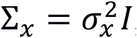, that is, each voxel is independent of the rest, then the Gaussian prior for the signal results is the usual Tikhonov regularization term. This optimization problem can be easily solved by steepest descent approach 
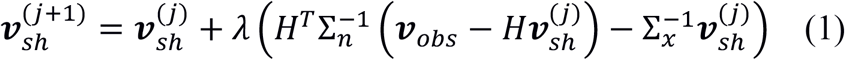
with ***v**_sh_*^(0)^ = ***v**_obs_*^26^. In the case of independent signal and noise voxels, 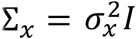 and 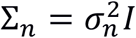, this iteration can be written as 
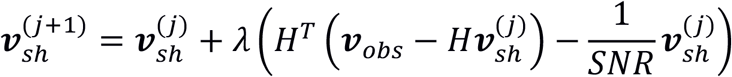
where the SNR is the signal to noise ratio defined as 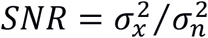. When the SNR is high, the deblurring term dominates, while for low SNR, the regularization term dominates. It can be easily shown that the standard Wiener solution 
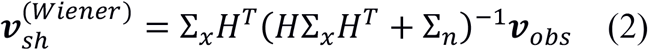
is a fixed point of our iterations. For proving this, let us define *D = H Σ_x_H^T^* + *Σ_n_* and study the gradient of the function *E*(***v***_*sh*_) at 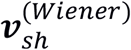 
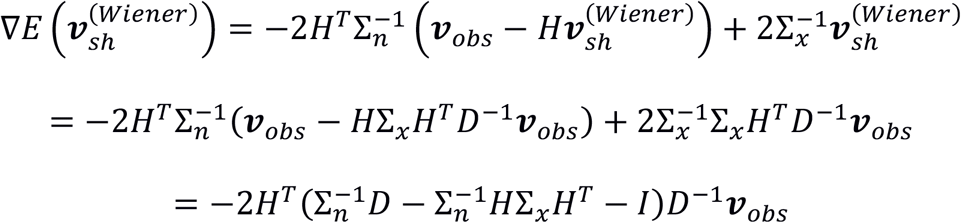

If we now substitute the definition of *D* in the parenthesis, we obtain 
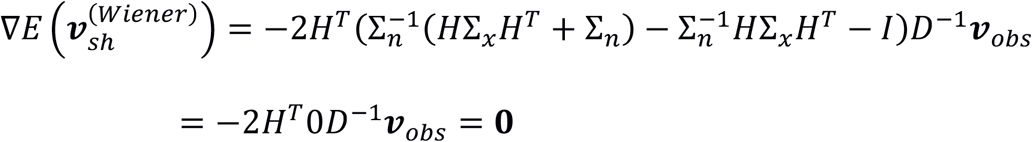

This small digression proves that our iterative algorithm in Eq. 1 converges to the Wiener solution in Eq. 2, with the advantage that it can be efficiently implemented in Fourier space; while Eq. 2 involves an inversion of a formidable matrix, which is computationally impractical. This iterative formula is repeated until a convergence criterion is reached. Our algorithm stops when the change between two successive iterations is less than 1%. The choice of the step size, *λ*, should be enough small to guarantee the convergence, but also large enough to speed-up convergence. We have observed that the following step size is a good compromise between both objectives: 
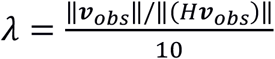

## Application of other sharpening methods

A number of sharpening methods, such as Relion post-processing^5^, Phenix AutoSharpen^6^ and LocScale^7^ were used for comparison purposes. AutoSharpen was applied with the default parameters, only taking into account the resolution reported in each case. For LocScale, each pdb was aligned to its corresponding map and applied as described previously^7^. For TRPV1, PDB ID:3j5p was used and for the 80S ribosome PDBs ID: 3j79, 3j7a were used.

## Code availability

The source code (LocalDeblur) can be found at https://github.com/I2PC/scipion/tree/release-1.2.1/software/em/xmipp and can be run using *Scipion* (http://scipion.cnb.csic.es) (release numbers greater than or equal 1.2.1).

## ACKNOWLEDGMENTS

The authors would like to acknowledge economical support from: the Comunidad de Madrid through grant CAM(S2017/BMD-3817), the Spanish Ministry of Economy and Competitiveness (BIO2016–76400-R), and the European Union and Horizon 2020 through grant INSTRUCT-ULTRA (INFRADEV-03–2016-2017, Proposal: 731005), iNEXT (INFRAIA-1–2014-2015, Proposal: 653706) and West-Life (EINFRA-2015–1, Proposal: 675858)

## AUTHOR CONTRIBUTIONS

J.M.C., C.O.S.S. and J. V. conceived the idea for this study. E.R.-A and J.L.V. wrote the code and the implementation in *Scipion*. P.C. and D.M. helped in the implementation in *Scipion*. E.R.-A designed the experiments and refined models. R.M. helped in the experiments and their interpretation. M.M. and R.M. helped in refined models. E.R.-A, J.L.V., J.M.C. and C.O.S.S. wrote the manuscript. All authors commented on and edited the manuscript.

